# Novel mechanistic insights for catalytic bioluminescence of mammalian Gaussia Luciferase through mutant and ancestral analysis

**DOI:** 10.1101/2025.06.16.659893

**Authors:** Raina Borum, Michael Lanzillotti, Aniruddha Sahasrabuddhe, John Hui, Victoria Cochran Xie, John Ferbas

## Abstract

A mechanistic basis for luciferase bioluminescence provides a glimpse into its evolutionary role for organism survival, as it provides a blueprint to engineer luciferase enzymes for advanced technological applications. *Gaussia* Luciferase is among the brightest natural luciferases, but (1) the evolutionary development of its luminescence behavior remains unclear, (2) recent fundamental studies utilized *E. Coli* expression systems instead of eukaryotic expression systems, and (3) notable mutants have been discovered but not integrated into comprehensive mechanistic analysis. We describe new mechanistic observations from GLuc by addressing these gaps. We monitored fluorescent coelenerazine-to-coelenteramide conversion to study turnover kinetics of mammalian-derived GLuc; this assay characterized the positive cooperativity kinetics of GLuc. The non-luminescent mutants, R76A and R147A, still turn over the substrate with high efficiency, each demonstrating sustained positive cooperativity. Through mass spectrometry, mutational analysis, and analytical liquid chromatography, we demonstrate that GLuc undergoes methionine oxidation during substrate turnover, and that this impacts the flash-type luminescence of the luciferase; we did not observe indications of covalent attachment with the substrate, product, or their intermediates. Chromatography on luciferases derived from ancestral sequence reconstruction highlighted that the extent of methionine-induced surface changes was greater for earlier ancestral luciferases. Ancestral sequence reconstruction also revealed that earlier ancestral copepod luciferases produced less light when compared to GLuc.

## Introduction

Bioluminescence has indispensable utility in nature and technology. Luciferases are bioluminescent enzymes that are used by an expansive range of animals; they are optically tuned for the functional phenotype, such as for inter-species communication, camouflage, mate attraction, or predator avoidance^2, 3^. Luciferase proteins are therefore quite diverse, with wildtype emission wavelengths ranging from blue to orange, luminescent half-lives ranging from seconds to minutes, and varying degrees of light intensities.

In the lab, luciferases are *in vitro* and *in vivo* reporters for diversified imaging studies and biochemical assays. Light from incident chemistry is more sensitive than light from incident light, making bioluminescence a more desirable reporter modality over fluorescence; bioluminescence is generally superior with wider dynamic ranges, lower background, and higher signal-to-noise (S/N) ratios^4^. The earliest application of luciferases was to monitor targeted protein expression *in vivo*^5^. They have been more recently developed to detect other molecular targets and events such as protein-protein interactions, protein degradation, protein secretion, reactive oxygen species generation, cell death, and so on^6, 7^. To advance their reporter efficacy and throughput, luciferases have been engineered for signal amplification, NIR-region wavelength emission, and increased sensitivity. They are currently relied upon for significant experiments involving target and drug discovery, drug efficacy, and high throughput screening^4, 8, 9^.

Gaussia Luciferase (GLuc) is an enzyme with exceptional technical potential. GLuc is derived from the marine copepod *Gaussia princeps* and is over 1000-fold more active than conventionally used sea pansy (*Renilla reniformis*) and firefly (*Photinus pyralis*) luciferases, but it is a fraction of their size at 18.2 kDa after signal cleavage^10^. It emits blue light with a peak wavelength between 470-485 nm and has flash-type photostability. The protein is also demonstrably splitable for protein complementation assays and consumes a common marine luciferin, coelenterazine (CTZ), as its luminescent substrate^2, 11^. Several functional mutations have been identified; these have enhanced photon yield, extended photostability, and enabled color shifting^9, 12-14^. However, most of these mutations have not been investigated further in relation to structure-function mechanisms.

Novelty around GLuc is also propelled by recent and continuously emerging data about its structure and catalytic mechanisms. Wu et al published the first NMR solution structure of GLuc in 2020, but their luciferase introduced two mutations to support bacterial expression of the enzyme^15^. In 2023, the binding domain between GLuc’s structure and an analog of the CTZ molecule was estimated via NMR, but the catalytic residues were not elucidated^16^. Dijkema et al demonstrated that the flash-type emission of GLuc is caused by compromised physical integrity of the enzyme when it consumes the substrate, leading them to describe the enzyme as ‘suicidal.’ In 2024, only 83.5% of the backbone atoms could be assigned to GLuc’s wildtype NMR solution structure, demonstrating that GLuc is a highly flexible, ‘less structured’ enzyme with largely disordered regions ^17, 18^. Markova et al characterized GLuc’s relative *Metridia* luciferases from the copepod *Metridinidae* family, and surveyed the environmental (temperature, salinity) and kinetic stability of copepod luciferases^19^. These findings suggest GLuc has mechanistic and structural features that haven’t been observed in other marine luciferases, such as *Renilla* and *Oplophorus* luciferases.

Yet the evolutionary development of GLuc’s luminescent behavior is practically unexplored. Analysis of ancestral proteins through sequence reconstruction can elucidate how mechanistic or structural changes manifest into resultant luminescence changes^20^. This can provide foundation for the evolutionary development of GLuc, but it can also identify regions on the luciferase with heightened potential to be engineered for functional features. We hypothesize that studying ancestral luciferases derived from GLuc and its homologs will offer new data that can clarify the core mechanics of GLuc, and how its luminescence might have naturally developed.

Moreover, the most recent and detailed fundamental studies on GLuc entirely relied on *E. Coli*-derived proteins. A persistent challenge in copepod luciferase studies has been the inability to reproduce results, such as luminescent intensity differences between mutants and the activity of fragmented GLuc. The results typically differ when the expression system (bacterial vs mammalian) differs^19^. GLuc is not only a secreted protein, but it contains ten Cysteines that all participate in disulfide bridging. Therefore, we hypothesize that a eukaryotic expression system would better accommodate a native folding pathway for the protein, making it a more accurately produced enzyme for downstream analysis.

Here we report mechanistic and evolutionary analysis of mammalian-derived GLuc. GLuc and all the studied mutants were expressed from human embryonic kidney cells. We explored hypothesis-driven single point mutations, developed a substrate turnover assay independent of luminescence, assessed protein changes through analytical mass spectroscopy and chromatography, and implemented mammalian derived ancestral luciferase analysis. Through this work, we propose a basis for the evolutionary development of GLuc’s bioluminescent behavior, we describe a newly observed mechanism that impacts its flash-type behavior, and we elucidate the role of certain residues regarding luminescent output versus substrate oxidation.

## Results

### Mutant screening: luminescence can be impacted throughout diverse locations of the sequence

We developed a computational method to approximate changes in physiochemical properties of proteins based on their amino acid sequence. Kim et al identified a region of GLuc where several one- and two-point mutations manifested in enhanced bioluminescent activity and spectral shifts through Kyte and Doolittle’s hydropathy mapping^12, 21^. We were interested in inspecting a wider set physiochemical properties of the sequence to (1) derive an impressionistic profile of localized physical potentials along the sequence, and (2) monitor how these properties change based on amino acid substitution. We chose physical properties from the AAindex such as flexibility, polarity, charge transfer and donor capabilities, net charge, accessible residues, buried volume, and Van der Waals volume to profile for GLuc, as we hypothesized these properties could provide insights onto the structural and dynamic capabilities of the enzyme^22^. To do this, we converted the string of amino acids into a property score *m* x *n* matrix, where *m* rows represent the positions along GLuc’s sequence, and *n* columns represent each property to be scored. A representative heatmap was derived from the normalized scores in the matrix **(Figure 1a)**. For simulated mutation prediction purposes and analysis, matrices of mutants could be subtracted from the wildtype matrix to inspect the resultant local changes from that mutation **(Figure S1a-b)**. This profiling was used to guide our choice in single point mutational screening.

**Figure 1.**
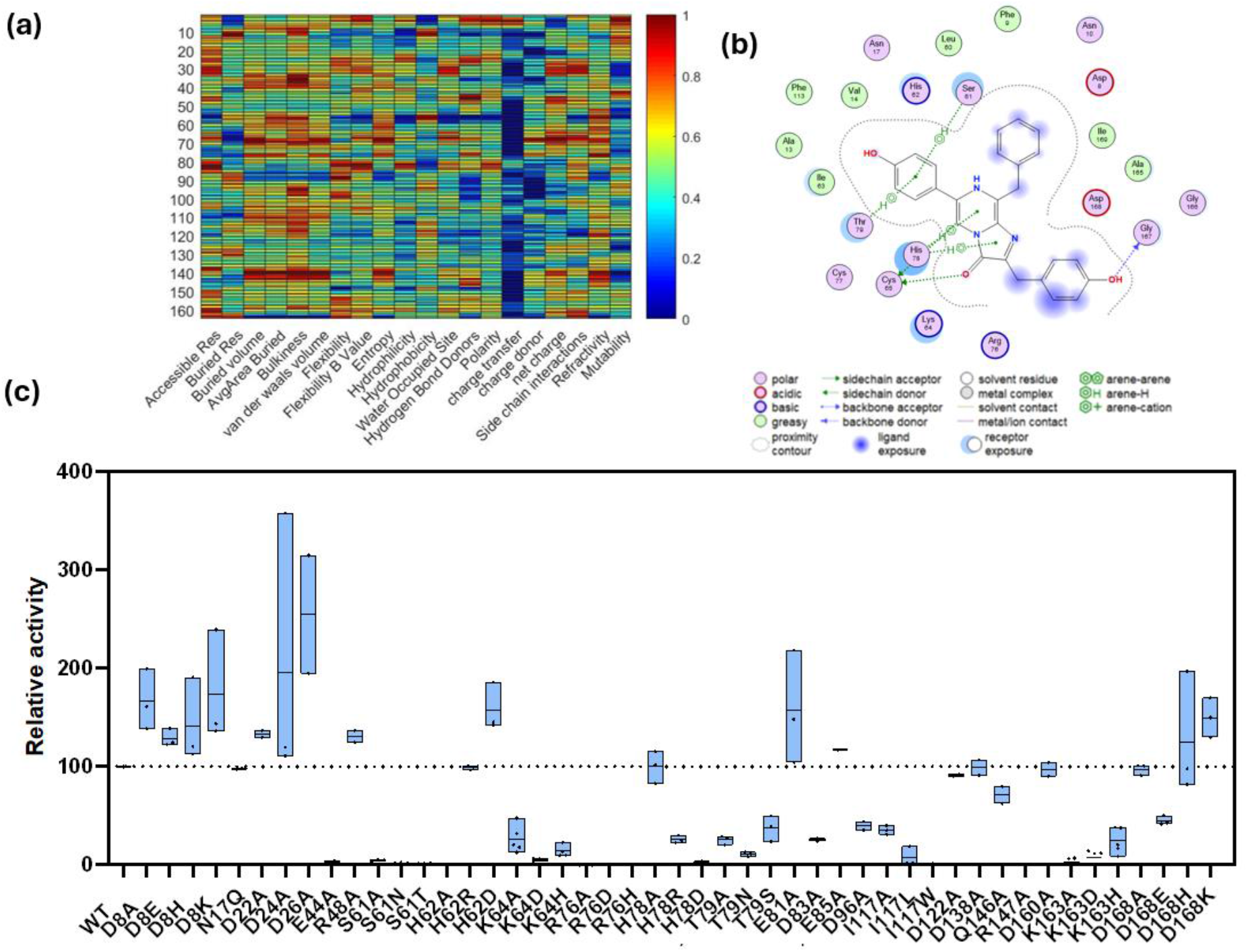
**(a)** Amino Acid-derived physio-chemical profile of Wildtype GLuc based on sequence. The color bar represents the normalized scores for the properties. **(b)** Molecular Docking illustrates the predicted binding domain of GLuc for coelenterazine. **(c)** Single mutation screening results show residues that have capacity to be mutated for enhanced bioluminescence (ex D8, D168), while some residues are more sensitive to mutations (ex: S61, I117)

We supplemented this computational analysis by employing Molecular Docking simulations between Wu et al’s solution structure (PDB 7D2O) and coelenterazine. From this, we compared the predicted binding domain residues with the local regions identified from the sequence profiling **(Figure 1b).** CTZ is nonpolar, so many hydrophobic residues were predicted to embody the binding pocket. Several acidic-based residue-CTZ interactions were also predicted.

Three regions gained outstanding interest after these analyses. The residues between positions 80 to 100 on the sequence were not only relatively hydrophilic, as first realized by Kim et al^12^, but their scores in flexibility, accessible residues, and charge transfer potential were high, with a local negative net charge. Besides this region, the first and last 10 residues of the sequence shared a similar profile. The molecular docking simulation also identified a binding domain that involves residues from all these regions **(Figure 1b)**. In these areas, we particularly analyzed charged residues to investigate their role in oxidation, hypothesizing that such flexible and accessible domains would accommodate active residues to process CTZ.

Single point mutations spanned from the 8^th^ to the final residue of the GLuc sequence **(Figure 1c).** Several anionic residues (D8, D24, D26, D168) yielded luciferases with enhanced bioluminescent activity when they were mutated. When D8 and D168 were substituted with cationic resides such as Histidine and Lysine, luminescent enhancement was still observed (**Figure 1c)**. Most notably, R76A and R147A each resulted in an entire turn-off of luminescent signal. Most substitutions resulted in blue emission within the wildtype emission range, but S61N most notably resulted in a 10 nm redshift, although all S61 mutations led to a significant decrease in activity **(Figure S2a-e)**. None of the Cysteines were challenged due to their putative role in the total structure of GLuc.

### GLuc’s positive cooperativity is conserved within certain non-luminescent mutants, R76A and R147A

When CTZ was mixed in DMSO for *in vitro* oxidation, a signature fluorescence signal at excitation 330 nm emission 415 nm emerged **(Figure 2a,b)**. We corroborated this by characterizing a CEI standard and noticed consistent fluorescent properties (**Figure S3a**) and therefore hypothesized that monitoring a reaction between GLuc and CTZ at this fluorescence window could monitor the accumulation of CEI product, and thus the turnover of substrate to product.

**Figure 2.**
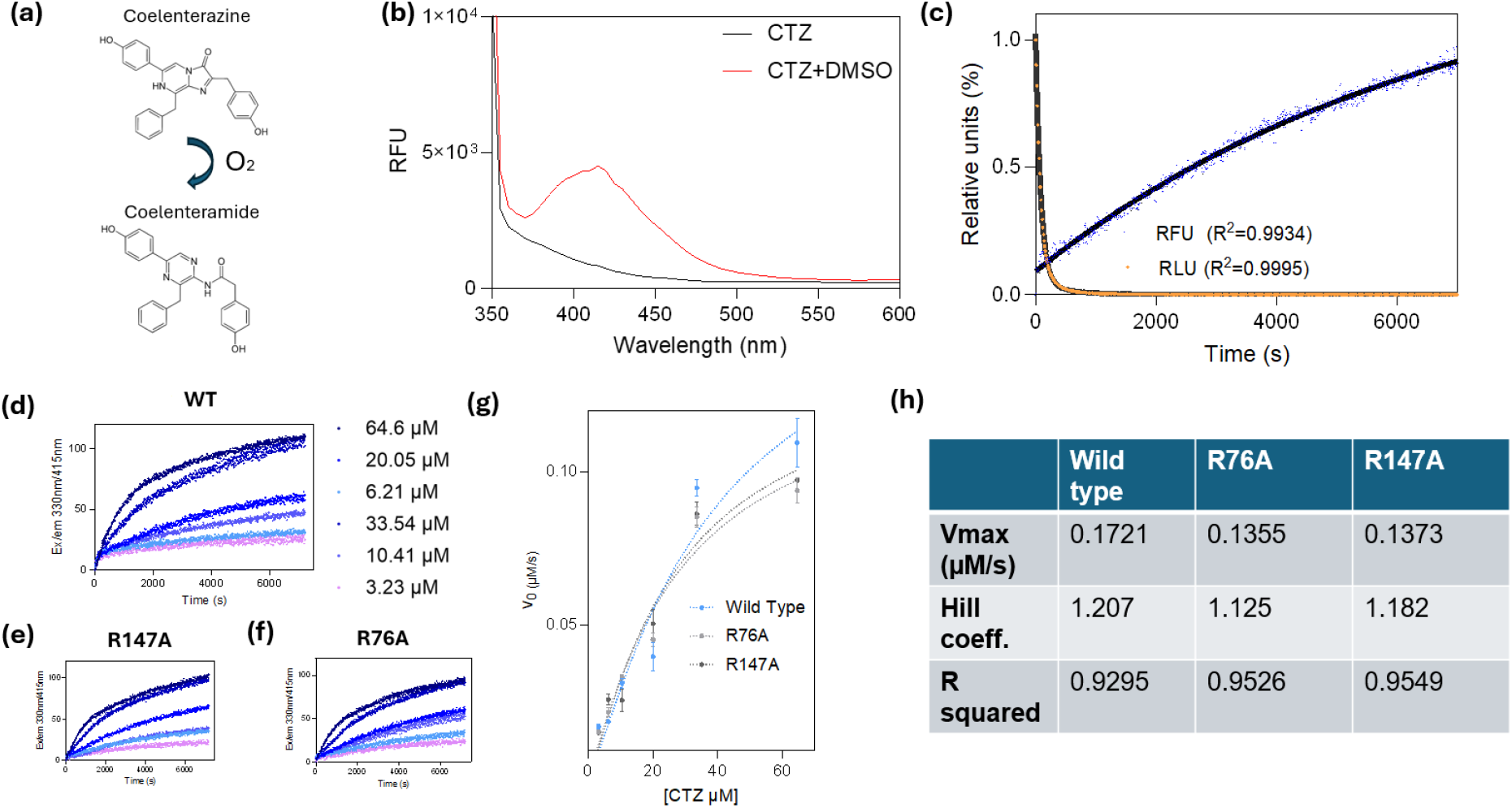
**(a)** Representative schematic of the transition from CTZ to CEI from oxidation. (**b)** Fluorescent emission spectra (ex 330 nm) of CTZ compared to oxidized CTZ in DMSO shows oxidized CTZ adopts a characteristic fluorescence not seen in fresh CTZ. **(c)** comparison of luminescence and CTZ-oxidation related fluorescence progress curves for wildtype GLuc, showing that the rate of CTZ turnover is slower than the luminescent output “flash” rate. **(d)** Resultant progress curves of substrate turnover with GLuc based on substrate concentration. **(e)** and **(f)** show that although R76A and R147A respectively do not produce light, they are still able to oxidize and turn over the substrate. **(g)** reaction kinetic curves for wild type and dark mutants all adopt signature sigmoidal cooperativity curves **(h)** summary of reaction kinetics show wildtype is faster and more cooperative in nature than the dark mutants

We monitored the fluorescent progress curves of GLuc reactions with CTZ and compared this rate with the luminescence output of the reaction. We observed that the best fit model for the luminescence emission was a double order exponential, similar to previous reports^17^. However, the fluorescence associated with CEI production accumulated at a slower rate (**Figures 2c, S3b**). The dark mutants (R76A and R147A) produced similar fluorescent progress curves (**Figure 2e-f**). We wanted to validate that the CEI accumulation from these mutants is caused by the intact enzyme rather than nonspecific oxidation; we did not see progress curves when we deactivated R76A, either with heat or acidic quenching, and any autooxidation of freshly prepared CTZ solution in PBS is relatively subtle in comparison (**Figure S4)**. These results support that these residues are utilized for light production rather than catalytic turnover.

When the assay is run under variable CTZ concentrations, the resultant progress curves can be used to approximate the substrate turnover kinetics of GLuc **(Figure 2d-f).** The kinetic profile illustrated a characteristic sigmoidal curve (**Figure 2g**). Like other reports on GLuc kinetics, we observed positive cooperativity, with a Hill coefficient of 1.21 (R2=0.93). Unexpectedly, R76A and R147A also showed some degree of positive cooperativity, with slightly lower Hill coefficients of 1.13 (R2=0.95) and 1.18 (R2=0.95) respectively **(Figure 2h).** However, the dark mutants had slower turnover velocities relative to the wild type **(Figure 2h).**

### Methionine Oxidation contributes to the flash-type emission of GLuc

Fresh GLuc replenishes luminescence of an ongoing reaction more efficiently than fresh CTZ, so we wanted to inspect post-reaction adducts and other potential covalent modifications. (**Figure S5a-b**) We therefore characterized the full intact protein mass of GLuc with LC-MS. Dijkema et al previously reported a 409.5 Da mass increase after GLuc reacts with CTZ^18^. We were unable to reproduce these results. We instead consistently observed abundant species of the protein sample with +16 and +32 Da shortly after the start of the reaction. (**Figure 3a,c, Figure S6**) To corroborate that these mass changes are reaction-induced rather than product-induced, we incubated GLuc with standard CEI. We did not observe any mass changes in this case **(Figure 3b)**.

**Figure 3.**
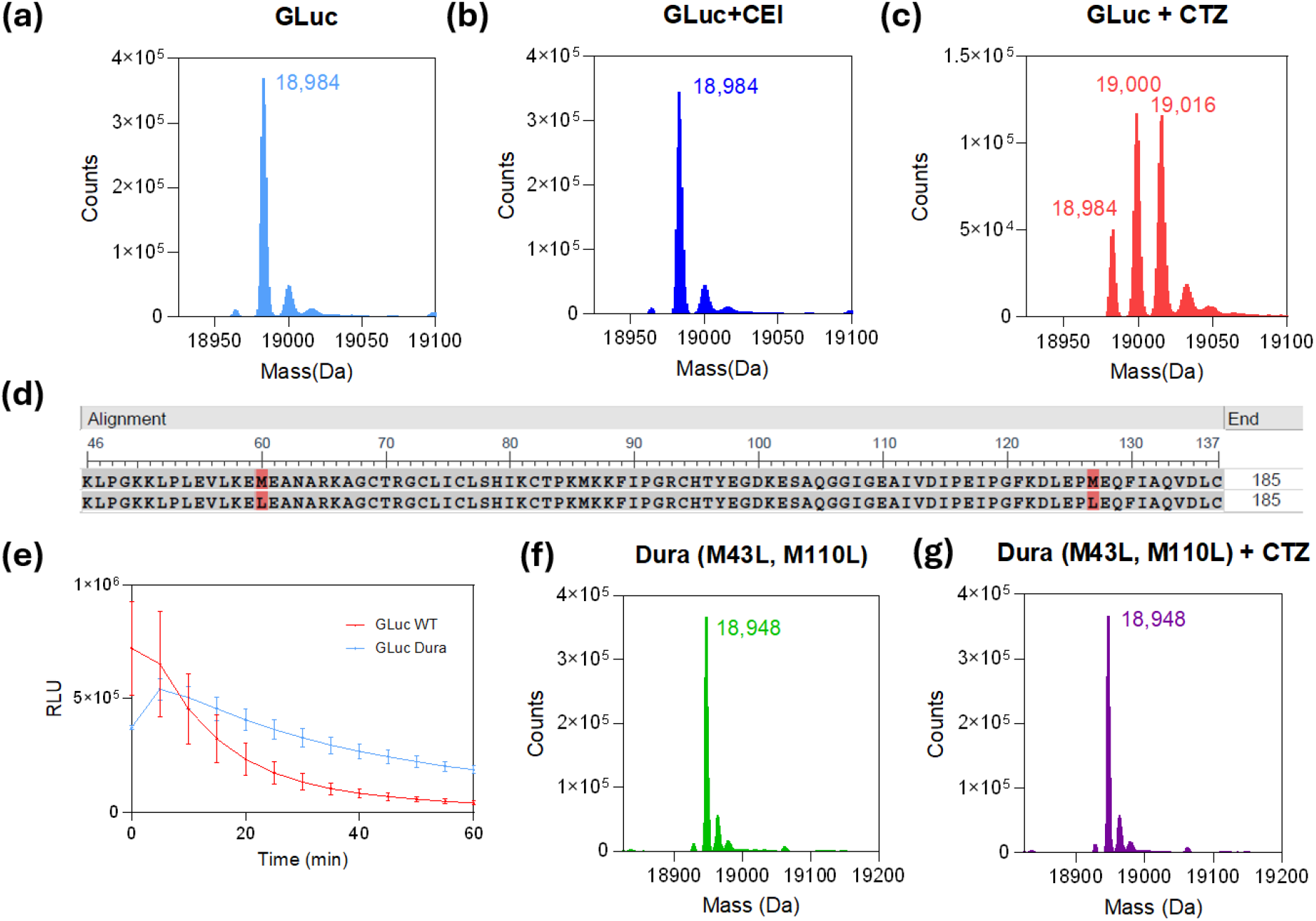
**(a)** Deconvoluted sample of GLuc reacted with CTZ, shows a +16 Da increase when compared to both (**b)** GLuc with CEI, the product, and **(c)** unreacted GLuc **(d)** photostability comparison between wildtype GLuc and Welsh et al’s GLuc Dura mutant, with the mutations highlighted. **(e) and (f)** show that even after incubation with CTZ, the total mass of GLuc Dura does not change.

We hypothesized that the 16 Da and 32 Da mass increases were caused by methionine oxidation^23^. The wild type has three methionine residues in its sequence, excluding the starting signal residue. Meanwhile, Welsh et al introduced a now-commercial variant of GLuc, ‘GLuc Dura,’ with two methionine mutations, M43L and M110L, that resulted in a glow-type emission (significantly prolonged bioluminescent half-life)^13^ **(Figure 3d-e)**. We tested this mutant similarly, and no mass increases were observed even 48 hours after the start of the reaction. **(Figure 3f-g).**

We wondered if other copepod luciferases share a similar mechanism when they react with CTZ. To explore this further, we tested the luciferase from the copepod *Pleuromamma Abdominalis* (PaLuc2), a relative of GLuc in the same *Metridinidae* family^24^. In this case, we also observed +16, +32, and +48 Da species. (**Figure S7a-b**). PaLuc2 and GLuc share 81.9% sequence similarity and are part of the same family.

This mass change was also studied with respect to reaction time points. Reactions were quenched both early and well into the initial rate period (1 min and 5 min), when the reaction was approaching saturation (15 min), and after saturation was reached (1 hour, 24 hours, 96 hours). We observed that the

+16 oxidized species emerged immediately, but after the reaction completed far after the start of the plateau (24 hours, 96 hours), a majority of the sample had +32 species with increasing presence of +48 species (**Figure S8a-g**)

### Disulfide bond mapping of mammalian GLuc

Putative disulfide bonds were identified based on the differences between the chromatograms collected of the reduced and nonreduced GLuc Lys-C digestions, shown in **Figure 4**. Three disulfide-bound peptides were identified by MS (including potential missed Lys cleavages, denoted by A’, A’’.), with the corresponding reduced peptides identified by MS in the reduced aliquot. These peptides accounted for five disulfide bonds including all 10 Cys residues present in the GLuc sequence. Disulfide bonds A and B correspond to peptides bound by a single disulfide; C65-C77 linking peptides L8 and L11,and C136-C148 linking peptides L14 and L16, respectively. Disulfide-bound peptide C contained 3 disulfide bonds based on its mass observed in MS1 (4237.947 Da), involving C52, C56, and C56 in peptide L7 and C120, 123, and 127 in peptide L13. The orientation of these disulfide bonds, however, is unclear based on MS1 data alone – either comprising 3 inter-peptide bonds or 1 inter- and 2 intra-peptide bonds.

**Figure 4.**
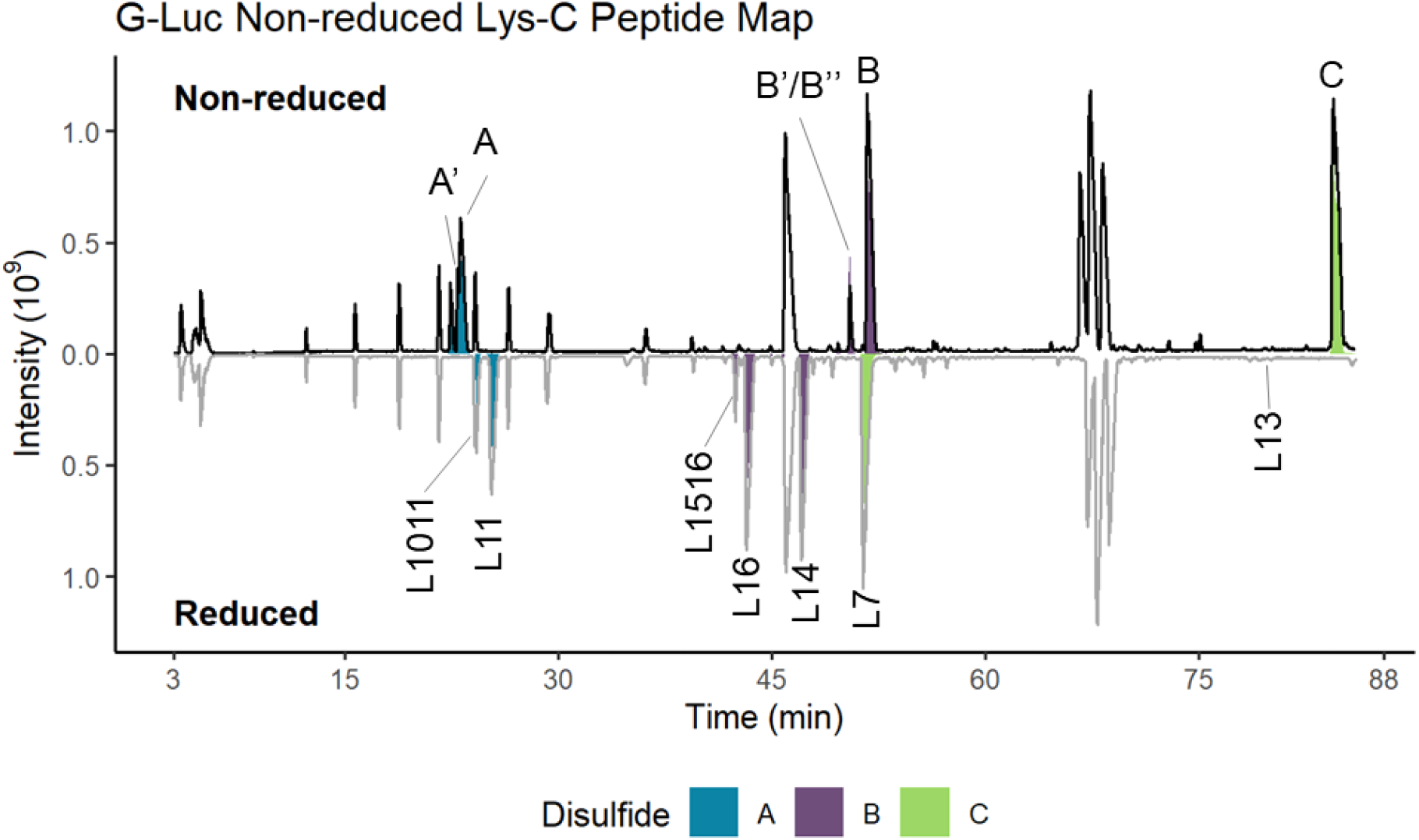
Annotated total ion current chromatogram for non-reduced (top) and reduced (bottom) Lys-C digests of GLuc for assessment of disulfide bonds. Extracted ion chromatograms of disulfide-bound peptides and their corresponding reduced peptides in are overlaid on the chromatograms and annotated.

MS/MS spectra of the 4+ charge state of disulfide-bound peptide C provides potential insights to the disulfide structure. HCD fragmentation of peptide L13 yielded high-abundance y-type fragment ions throughout the peptide backbone, up until C120 – consistent with the presence of an interchain disulfide bond at or after that residue. Additional fragment ions identified throughout the peptide backbone suggest that C120 contains the inter-chain disulfide bond and that C123 and C127 are bound in an intra-peptide bond.

### Ancestral Sequence Reconstruction and Characterization

We used ancestral sequence reconstruction to characterize the evolutionary development of the luciferase and its current luminescent behavior. We studied five ancestral sequences derived from FireProtASR tool^1^ (**Figure 5a, Table S1**). The root node ancestral luciferase (ANC26) had the lowest luminescent output, while GLuc remained the brightest luciferase amongst all ancestrals (**Figure 5b**). The photon yield of ancestral luciferases progressed into a comparable yield to GLuc with increasing proximity to GLuc. We could not find any correlation between kinetics and ancestral luciferases relative to GLuc, although we observed that while ANC 26 had the lowest relative velocity of the ancestrals, it had the highest Hill coefficient of 1.6 (R2=0.97). All ancestrals similarly emitted blue light like Gluc, with ANC31, ANC35, and ANC36 producing slightly blue-shifted peaks between 464-466nm. We could also not find a correlation with bioluminescent half-life, although ANC 31,35,and 36 showed a more glow-type emission (**Figure 5c-e**).

**Figure 5.**
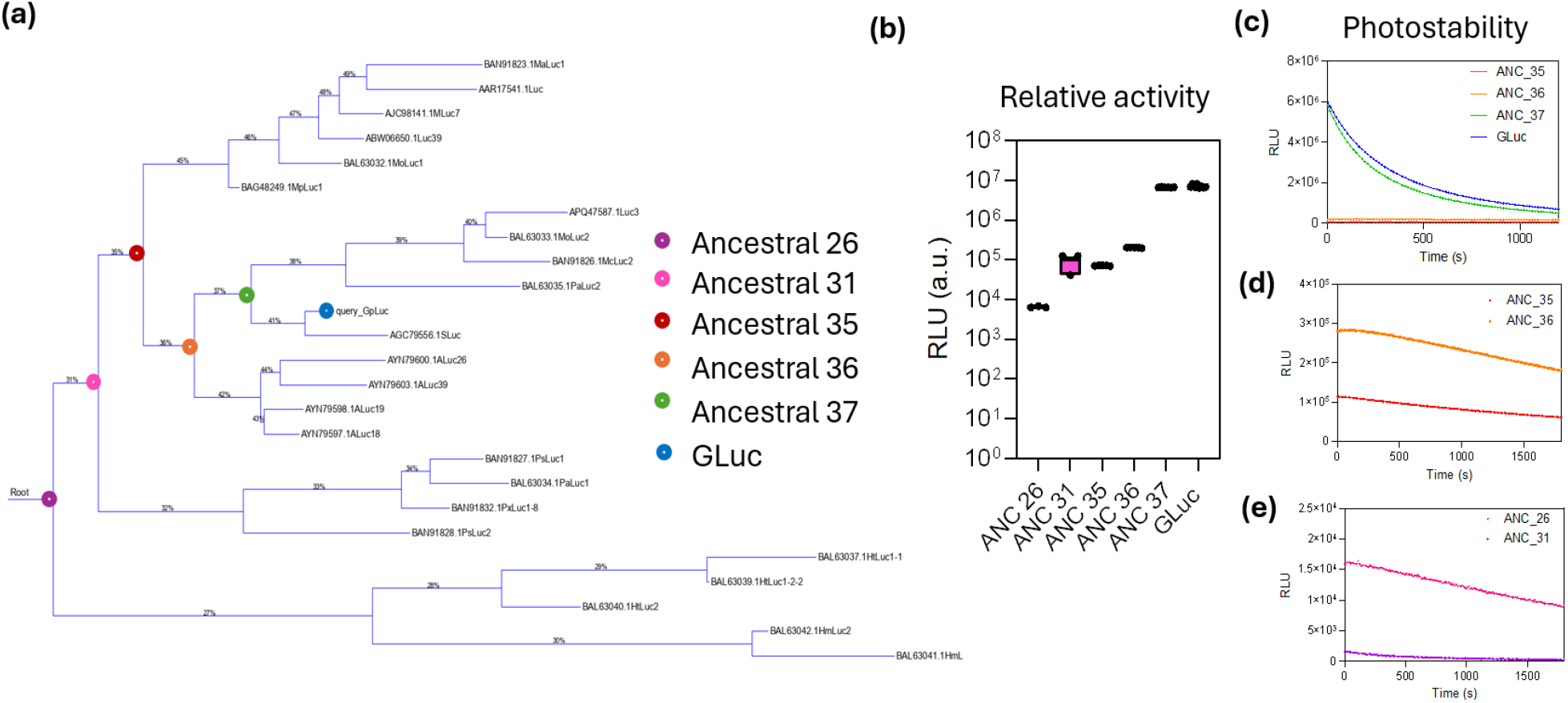
**(a)** Resultant ancestral tree derived from Ancestral Sequence Reconstruction^1^. The analyzed ancestral luciferases are highlighted by the colored circles; these sequences were mammalian expressed and then studied for bioluminescence, as seen in **(b),** where the ancestrals developed increased luminescent activity as they approach from the root node to GLuc, and **(c)** shows varying photostability with **(d-e)** providing zoomed curves.

Like GLuc, the ancestrals produced species with +16 Da, +32 Da, as Methionines were conserved. When we analyzed reacted and unreacted GLuc and ancestral luciferases through analytical RP-HPLC, we were able to separate these oxidized species (**Figure 6b, S11-12**). After resolving these species on RP-HPLC, we compared the retention time differences between the native and oxidized protein species, between GLuc and the ancestral sequences. In the order of the root node to GLuc, the ancestral luciferases were in order of decreasing retention times between their unoxidized and +16 Da species (**Figure 6c)**.

**Figure 6.**
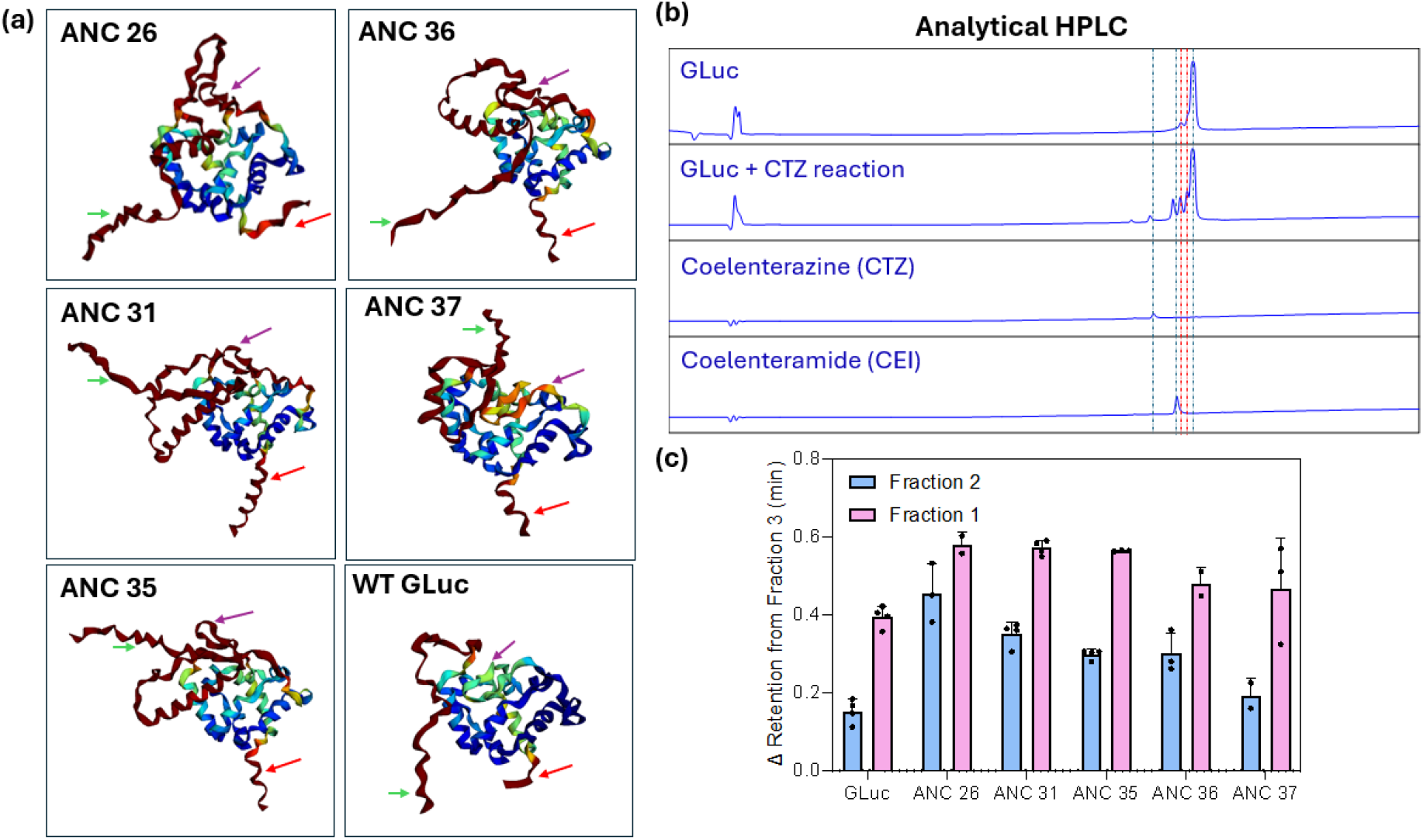
**(a)** Resultant AlphaFold2 structural predictions of ancestral as well as Wildtype GLuc. The N-terminus and C-terminus are identified with the green and red arrows respectively, and the conserved flap motif is also identified with the purple arrow. The color scale ranks from low to high plDTT scores from blue to red, showing that the flap region for WT Gluc had the lowest relative plDTT score with respect to its ancestrals. Other angles of the ancestral luciferase structure predictions are shown in Figures S11-16. (b) representative RP-HPLC chromatograms reveal additional eluents of oxidized species (as indicated with the red dashed line) when GLuc reacts with CTZ. The blue dashed lines indicate the standard eluent fractions of GLuc, CTZ, and CEI. **(c)** Analytical RP-HPLC shows that the retention time between +16Da and +32 Da oxidized (Fractions 1 and 2) and non-oxidized species (Fraction 3) is larger for the ancestrals than for GLuc, suggesting significant structural changes from oxidation.

These results motivated us to compare predicted structures of the ancestral luciferases with GLuc. When we analyzed the ancestral luciferases via AlphaFold2, both similarities and differences between GLuc and all ancestrals were outstanding. All luciferases shared a similar core structure, with alpha helical bundles bound together by the conserved cysteines in the sequence. Additionally, all luciferases had a highly flexible “flap” motif, but the local plDTT scores for this flap were the lowest for GLuc. Finally, the ancestrals had longer N- and C-terminal tails with higher plDTT scores (**Figure 6a, S13-18)**.

## Discussion

Coelentertazine is turned over by *Gaussia, Renilla*, and *Oplophorus* (the NanoLuc wildtype) luciferases, as well as *Aequorin* and Obelins for light. However, all CTZ-consuming luciferases are different in structure and size, and the putative catalytic residue ensembles and dynamics of *Renilla* and Nano luciferases are not similar^25, 26^. The oxidation of CTZ can be accomplished by a variety of charged-propelled interactions. Mutations from the 8^th^ residue to the final residue of GLuc manifested into luminescence enhancement, supporting the postulate that GLuc uses its entire body—both core and intrinsically disordered regions—to process CTZ for efficient light production. This additionally supports GLuc’s need to be relatively less structured than the *Renilla* and Nano luciferases.

The chemical transition from coelenterazine to coelenteramide can occur radiatively or nonradiatively; while nonradiative decay (mechanical energy or heat dissipation) is typically more energetically favorable, radiative decay is necessary for photon emission. CTZ is thus still capable of oxidizing into the CEI product molecule without emitting light. We therefore hypothesized that establishing a relationship between the bioluminescence kinetics and the substrate turnover would not directly probe the chemical catalysis kinetics of GLuc. The fluorescence association CEI assay confirmed that GLuc was still turning over CTZ with either R76A or R147A mutations; these mutants thus indicate the role of R76 and R147 for radiative emission. The sustained positive cooperativity in these mutants suggest GLuc’s dynamics make it a highly efficient enzyme. This may also suggest a possibility that luminescence may have been incorporated only after the basic protein evolved with inherent positive cooperativity. It is therefore likely that the Arginine residues use specific binding interactions to configure CTZ into the catalytic domain with minimized remaining flexibility for nonradiative decay. We agree with Dijkema et al that the guanidium stacking mechanism is likely responsible for the rigid anchorage of CTZ within the binding pocket, and thus the radiative decay^18^.

Our intact mass spectrometry results differ from Dijkema et al’s, allowing us to draw conflicting conclusions about the effect of GLuc’s structure on its flash-type behavior. Our inability to reproduce their results may be in part caused by our use of different expression systems; this has already been a demonstrably prominent discrepancy in copepod luciferase studies. While *E. Coli* expression is essential for isotope labeling and subsequent NMR characterization—the only viable method to chart the structure of a small and flexible protein like GLuc—we believe that a mammalian expression system is more suitable for foundational and consistent catalytic studies of these luciferases, where high resolution structural analysis is not required. Moreover, our observations of methionine oxidation not only remain consistent with PaLuc2, as well as GLuc’s derived ancestral sequences, but it can be readily explainable by the putative oxidative catalysis GLuc and other coelenterazine-consuming luciferases employ.

Many marine luciferases are largely categorized as oxidoreductases and catalyze substrate turnover through the oxidation of coelenterazine. A key effect from oxidoreductase activity is the significant generation of reactive oxygen species^27^. Methionine oxidation is often regarded as a structurally protective mechanism against the oxidation of aromatic residues like Tryptophan and Phenylalanine^28^. It is therefore reasonable to suggest that GLuc, an oxidoreductase, would employ Methionine oxidation as a protective mechanism for its structure for effective catalysis. Nonetheless, it has been shown that the structure or surface of the protein is still impacted after Methionine oxidation; the GLuc separations on RP-HPLC suggest that there is considerable surface change after the reaction.

The simulated physiochemical profile heatmap of GLuc in **Figure 1a** illustrates how M43 is in a highly accessible region on the sequence, and Welsh et al corroborated through single point mutation analysis that M43 led to the most impactful change in glow-type emission when compared with M69 and M110^13^. The Dura mutant’s absence of these oxidized species further suggest methionine oxidation has some contribution to the protein’s dynamics that modulate its luminescence half-life. Previous reports have demonstrated the catalytic relevance of certain methionine oxidation events. For example, Gan et al showed that for 15-lipoxygenases, the oxidation of active-site Methionine also led to enzymatic inactivation, however, the substitution to Leucine maintained similar auto-inactivation, suggesting the methionine was a protective antioxidant rather than part of a catalytic residue ensemble^29^. Methionine oxidation seemed to persist with ancestral luciferases and *Pleuromamma abdominalis* as well, validating that the conserved methionines have a critical contribution to the bioluminescent emission phentotypic trait for copepod luciferases.

The unambiguous assignment of the C136-C148, and C77-C65 pairs from LysC digestion, combined with their unambiguous assignment along with C59-C56 pair from Dijkema et al’s tryptic digestion solidify these three putative pairs for GLuc’s higher order structural configuration. The remaining pair configuration remains ambiguous, although all ten cysteines clearly participate in disulfide bridging. These results perpetuate enigma in the dynamics or diversity of GLuc’s structure; it is certainly possible that the remaining two cysteine pairs can both exist.

Ancestral sequence reconstruction offered critical perspective on how this luciferase might have been naturally selected. The AlphaFold results for the ancestrals shared some predicted features with GLuc; the flexible flap region, and a core structure with two pairs of helical bundles. The flap region of GLuc corresponds to a putative active site identified by Kim et al and corroborated by our physiochemical profiling^12^. Moreover, several differences were interesting: ancestral sequences closest to the root node had increasingly dense regimes with high plDTT scores. The oxidized species are heavier than the native protein structure, and the ancestral luciferase oxidized species eluted out earlier than the native protein, relative to the elution times between the oxidized and native species of GLuc. These results might imply that due to the relatively lower estimated structural rigidity of the ancestral luciferases, their surfaces have a more pronounced change after their reaction with the substrate, rendering them less stable. This can provide insights as to how GLuc evolved and was selected for its ultimate phenotypic luminescence.

Bioluminescent copepods are ubiquitous and live in every ocean. *Gaussia princeps,* the organism that expresses GLuc, like copepods from the genus *Metridia* and *Pleuromamma* in the *Metridinidae* family, uses bioluminescence as a “distract or blind” mechanism for its predators. In a previous study by Takenaka et al, *Metridia pacifica* and *Pleuromamma* were compared alongside other copepod luciferases, but their luciferases produced the brightest luminescence, postulating that their specific “distract or blind” function favors brighter luminescent activity against predators^24^. The gradual increase in luminescence with each progressive ancestral luciferase sequence from the root node to Gluc supports Takenaka et al’s postulate, demonstrating that copepod luciferases evolved to become increasingly brighter with a more pronounced flash effect. Meanwhile, the gradual decrease in retention times between oxidized and unreacted luciferase species with each progressive ancestral sequence supports its evolution towards a more stable and compact structure for photonic yield.

## Conclusions

Here we introduced new mechanistic insights of mammalian expressed Wildtype GLuc. Our CTZ-CEI fluorescent assay confirmed that even non-luminescent GLuc can turn over CTZ with positive cooperativity. We discovered the presence of methionine oxidation on GLuc when it consumes its luciferin substrate CTZ, and observed its role in modulating the flash type emission. We employed ancestral luciferase analysis to observe how this conserved oxidation impacts the reacted species through analytical chromatography, and outlined the evolutionary development of GLuc’s bioluminescent activity through these ancestrals.

These insights can be used as potential transfer learnings to semi-rationally evolve GLuc into a functional bioluminescent probe. The luminescent Arginines and compact stability of the protein core body, for example, can serve as constraint features; the sequence along these regions may be conserved to maintain structural stability as well as luminescence ability. The residues immediately surrounding the oxidation site, the Methionines responsible for oxidation, the flap region, as well as the flexible N- and C-terminal tails, however, have potential to serve as mutationally rich regions that could be further evolved for modified and potentially optimized luminescence.

Collectively, these results demonstrate that copepod bioluminescence relies on enzyme dynamics that are practically unique from other luciferase species, including marine luciferases. They provide a new outline into how luciferases utilize a set of biochemical and intramolecular interactions to process the luciferin for light, how these interactions have critical relevance to an evolved and functional phenotypic trait, and a novel blueprint for engineered bioluminescence.

## Methods

### Computational Physoichemical Profiling

Physiochemical profiling was implemented via MATLAB. In summary, the input amino acid sequence (GLuc WT) was converted as a string into numerical scores based on reported property scoring in the AA index. This scoring method was based on Kyte and Doolittle’s sequence-score averaging technique, with a sliding window of 9^21^. The signal peptide was excluded from this analysis.

### Molecular docking

Molecular Operating Environment ‘MOE’ (Chemical Computing Group) was used to predict docking interactions between CTZ and wildtype GLuc. The structure used for GLuc (PDB 7D2O) was first added into MOE and quick prepped to an RMS gradient of 0.01 kcal/mol/A^2^. The Coelenterazine substrate molecule in the simulation was derived using the SMILES for the molecule, and it was washed and quick prepped before docking simulations. We used induced fit for method refinement, with five poses predicted. Ligand interactions were analyzed and used to derive our mutational screening library.

### Luciferase mutant activity screening

Mutant activity screening was carried out using Phosphine Buffered Saline (PBS) with 0.1% Tween-20 (PBST). Luminescence characterization was done with a SpectraMax M3 (Molecular Devices) plate reader, using 500 ms^-1^ integration time and half area white 96 well plates (Corning). All mutants were calibrated to 10 nM concentration in PBST and were plated. 100 µL of fresh 10 µg CTZ in PBST was then added to each well and the endpoint luminescence plate measurement was immediately taken. For photostability measurements, luminescence was measured at 500 ms integration for 10 minutes, with 2 second intervals. Luciferase mutants and wildtype GLuc were expressed, purified, and supplied through the HEK HTP service provided by GenScript, and they each had a 6xHis tag at the C-terminal of the sequence.

### Enzyme kinetic measurements

Wildtype and mutant GLuc was prepared and plated into black 96 well plates in an identical manner to the mutant library screening. Six different concentrations of fresh CTZ were prepared via half-log serial dilution in PBST, in triplicate. CTZ preparations were then added to the luciferase plate using a multichannel pipette. The reaction was subsequently read for fluorescence endpoint readings, using a 330 nm/415 nm excitation/emission window. The reaction was read through one hour, with 2 second kinetic measurement intervals. Readings were all done with the SpectraMax M3 platereader.

Kinetic calculations were derived using GraphPad Prism. Standard coelenteramide (CEI) (NanoLight Technologies) was used for a calibration curve at the same fluorescence wavelengths, and the slope was then used to calculate product conversion approximations for the monitored GLuc reactions.

### LC-MS for intact protein mass

Intact Protein Mass spectrometry was conducted on an Agilent 6200 TOF LC-MS system. To study post-reaction modifications on the intact protein, 10 µg of GLuc was reacted with 10 µg of CTZ in PBS; no tween or detergents were included. The reacted samples were sealed and left in the dark between 20 minutes to 3 days. Reacted samples, luciferase alone, substrate alone, and product alone were then diluted 2X in 0.1% TFA and analyzed. Data was collected and analyzed via Bioconfirm software and presented on GraphPad Prism.

### Kinetic-Oxidation species assay

For this assay, GLuc reacted with CTZ in PBS in a 96 well plate, without any surfactants. The plate was scanned at the Spectramax M3 plate reader at similar fluorescence reading conditions described in enzyme kinetic measurements. At the prescribed time points, plate scanning was paused, the plate was removed from the instrument, and the reaction wells for the given time point were quenched by adding TFA for a final concentration of 0.5% TFA. The plate was then returned to the plate reader and data was appended to continue the kinetic scan of ongoing, unquenched reactions. Inactivation of quenched wells was also validated by confirming no further growth of fluorescent signal. All samples were then analyzed for protein intact mass spectrometry in the Agilent 6200 LC-MS system.

### Non-reduced Peptide Map

Disulfide structure was characterized by peptide map, involving a bottom-up digestion of GLuc under non-reducing conditions. GLuc samples were denatured in a guanidine hydrochloride (Thermo Scientific) solution with addition of N-ethyl maleimide (Sigma Aldrich) to modify any potential free Cys residues. After denaturation, the sample was diluted in Tris hydrochloride buffer, pH 8.0 (Invitrogen) to a final concentration of 0.5 mg/mL and digested for 1 hour at 37C with Lys-C (Wako) at an enzyme to protein ratio of approximately 1:10. A subaliquot of the digest was reduced with TCEP. Both the reduced and nonreduced aliquots were quenched to a final concentration of 0.2% trifluoroacetic acid.

LC-MS analysis was conducted on an Orbitrap Q-Exactive Plus mass spectrometer equipped with the BioPharma option coupled to a Waters Acquity H-Class liquid chromatograph. Mobile phase A consisted of 0.1% trifluoroacetic acid in LCMS grade water (Fisher Chemical), and mobile phase B consisted of 0.1 trifluoroacetic acid in LCMS grade acetonitrile (Fisher Chemical). The protein digests were injected onto a Waters Acquity C4 column (1.7 micron, 150×2.1 mm) and separated using a 0-37%B gradient. A top 5 MS/MS method was employed, utilizing an MS1 resolution of 60,000, and MS/MS resolution of 30,000, and a higher-energy collisional dissociation (HCD) activation of 27 NCE.

LC-MS data was analyzed by Mass Analyzer for disulfide bond identification. Disulfide bond identities were confirmed manually and visualized using R.

### Top-Down Mass Spectrometry

GLuc was prepared for top-down mass spectrometry by buffer exchange via 10k MWCO (Amicon) into water. The buffer-exchanged protein was diluted to an approximate concentration of 4 µM in a 1:1:2 solution of water, acetonitrile, and methanol, respectively, acidified with 0.1% formic acid. This solution was introduced to an Orbitrap Fusion Lumos mass spectrometer via heated electrospray ionization (HESI) with an ionization voltage of 3.5 kV. Sample aliquots were reduced during infusion by addition of TCEP to a final concentration of 40 micromolar, corresponding to an approximately 2X molar excess relative to the number of disulfide bonds.

The 13+ charge state of GLuc was isolated using the quadrupole (5 m/z width) and activated with HCD (32 NCE), electron-transfer dissociation with supplemental collision dissociation (EThcD, reagent ion AGC target of 7e5, reaction time of 10 ms, and 15 NCE of supplemental activation energy), and 213 nm ultraviolet photodissociation (UVPD, activation time of 50 ms or 125 pulses).

The top-down spectra were analyzed and visualized using R, and fragment identifications used for disulfide bond characterization were validated manually.

### Analytical RP-HPLC

RP-HPLC was completed on an Agilent 1200 Infinity system, with a Jupiter Column (150 × 2 mm, 5 micron, 300 Angstrom C4). 10 µg of each luciferase studied was prepared. Proteins were either analyzed alone or after they reacted with 5-10 µg of CTZ for 30 minutes. No tween or surfactants were used. All samples were then diluted 2X with 0.1% Trifluoroacetic Acid (TFA) in HPLC grade pure water before they were analyzed.

The RP-HPLC method was the same for all HPLC experiments: the mobile phase (0.1% TFA, MeCN) was brought from 5% to 85% in 18 minutes, raised to 100% after an additional 2 minutes, and lowered to 5% after an additional 3 minutes. Sample was collected at the apex of each eluent peak. Before further analysis, eluents were immediately buffer exchanged with a 7000 MWCO to PBS. Eluents were then either studied for intact mass measurements though LC-MS, or used to study recovered luminescence.

## Acknowledgements

We are deeply grateful to Iain Campuzano and Xiaomin Chen for the helpful and productive discussions on this work.

