## Supplemental Data for "Novel mechanistic insights for catalytic bioluminescence of mammalian Gaussia Luciferase through mutant and ancestral analysis"

Internal Use Only Ethics and Compliance

| Name | AA Sequence |
|---|---|
| ANC 26 | MMRLLSLVLFALICFYLIQAWPAEDEEDDDIIDIVGVEGKFGTTDLETDLFTLWKDYWGIGVDNDANNEKDVSGDRGRGGKL<br>PKKLSLEVLKEMEANARRAGCTRGCLIGLSKIKCTPKMKKFLPGRCHSYAGDPATGQGPLGGAIVDIPEIPGYKDLTPMEQFI<br>AQVDLCADCSTGCLKGLANVKCSDLLKKWLPTRCATFATQIQSEVDNIKGLGGDRDPPIG |
| ANC 31 | MPRGMMEILSKVLFALICFALVQANSQKLLPTENENKDDIIDIVGVEGKFGTETDLETDLFTIWEDMNVISRDTDVDANRANNT<br>NLVNGDRGRGKLPGKKLPLEVLKEMEANARRAGCTRGCLICLSKIKCTAKMKKYIPGRCHSYEGDKETGQGGIGGAIVDET<br>VDIPEIPGFKDLEPMEQFIAQVDLCADCTTGCLKGLANVKCSDLLKKWLPDRCASFANKIQSEGEVDNIKGLAGDRDELIGIK<br>QTDKGK |
| ANC 35 | MEILSKVLFALICFALVQANPTENKDDIIDIVGVEGKFGTTDLETDLFTIWEDMNVISRANNTNLVNGDRGRGKLPGKKLPLEV<br>LKEMEANARRAGCTRGCLICLSKIKCTAKMKKYIPGRCHSYEGDKETGQGGIGGAIVEIPGFKDLEPMEQFIAQVDLCADCT<br>TGCLKGLANVKCSDLLKKWLPDRCASFADKIQSEVDNIKGLAGDRDELIG |
| ANC 36 | MGILSKVLFALICFALVQANPTENKDDIIDIVGVEGKFGTTDLETDLFTIWEDMNVISRANRANNDNGDRGRGKLPGKKLPLEV<br>LKEMEANARRAGCTRGCLICLSHIKCTAKMKKFIPGRCHSYEGDKETGQGGIGEAIVEIPGFKDLEPMEQFIAQVDLCADCT<br>TGCLKGLANVKCSDLLKKWLPSRCATFASKIQSQVDKIKGLAGDRDELIG |
| ANC 37 | MGILSKVLFALICIAVVQAKPTENNEDIIDIVAVAGNFATTDLANRANNDGKLPGKKLPLEVLKEMEANARRAGCTRGCLICLSH<br>IKCTAKMKKFIPGRCHSYEGDKESAQGGIGEAIVVDIPEIPGFKDLEPMEQFIAQVDLCADCTTGCLKGLANVQCSDLLKKWL<br>PQRCATFASKIQSQVDKIKGLAGDRDELIG |

**Table S1** AA sequences of the ancestral luciferases studied, from N to C terminus. The conserved methionines are highlighted in yellow, while the conserved arginines are highlighted in green.

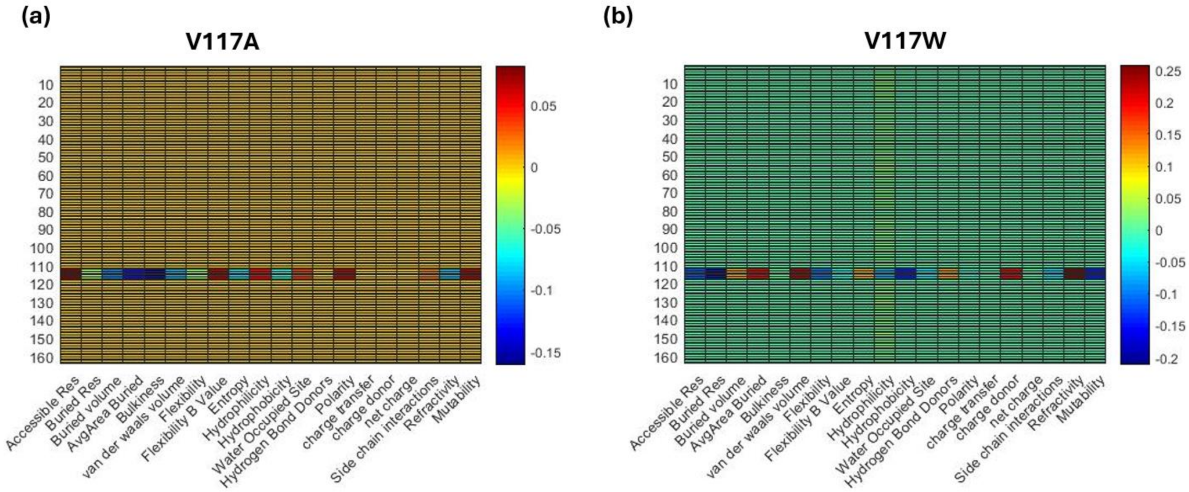


**Figure S1**. Subtractive heat maps between the mutant of interest with the wildtype GLuc shows a representative approximation on how amino acid substitutions can manifest in a different set of physiochemical property score changes.

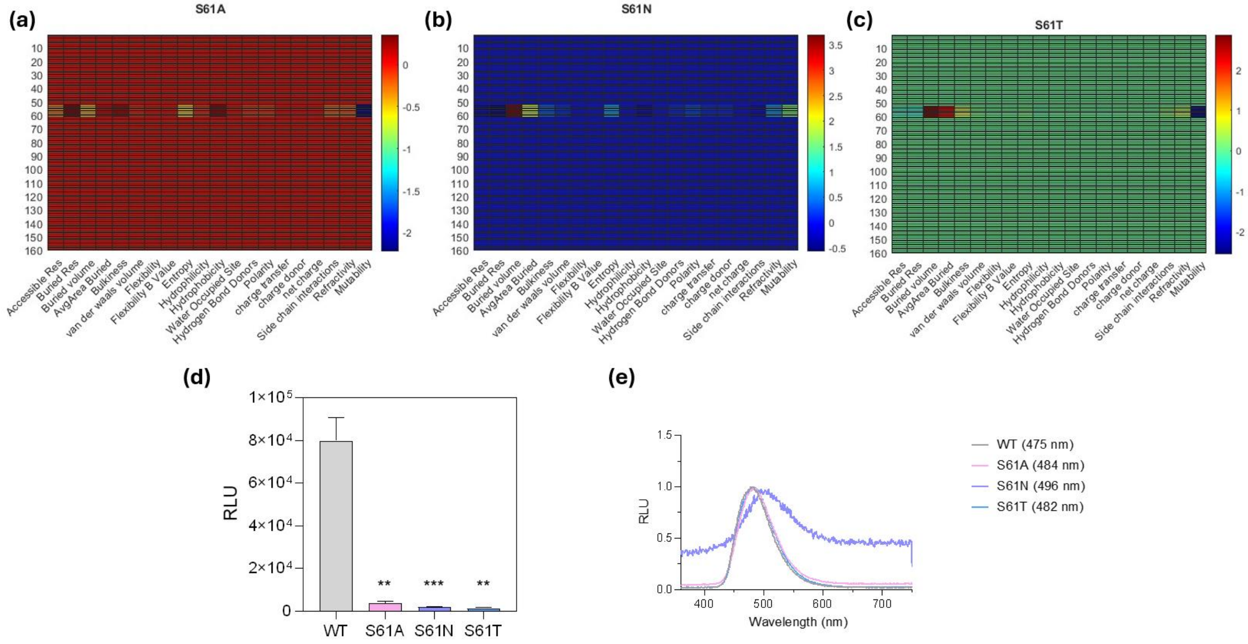


**Figure S2**. Representative characterization of S61 mutants. (a-c) show the possible changes from the three mutants that were inspected. Although each mutant lost significant activity as seen in (d), S61N resulted in a redshifted mutant with an emission spectral peak at 496 nm.

(a)

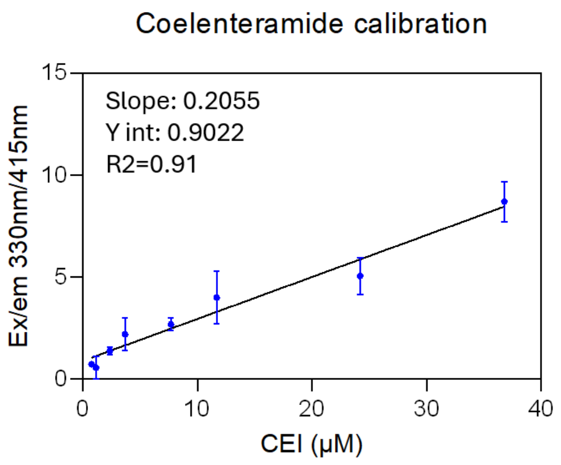


(b)

| Measurement | Best fit model |
|---|---|
| Luminescence decay | $\mathrm{F}(t) = f(0) + a(1 - e^{-k_1 t})+\mathrm{b}(1 - e^{-k_2 t})$ |
| Fluorescence association | $\mathrm{F}(t) = f(0) + \big(a - f(0)\big)(1 - e^{-k_1 t})$ |
| Dijkema et al, Protein Science doi: 10.1002/pro.4023 | $\mathrm{F}(t) = f(0) + a(b - (ce^{-k_1 t}+(1 - c)e^{-k_2 t})$ |

**Figure S3**. (a) Calibration curve of a Coelenteramide standard (NanoLight Technologies) fluorescence intensity in PBS using a half log dilution. (b) charted fit models corresponding to the luminescence vs fluorescence progress curves outlined in Figure 2c in the main manuscript show the CTZ-CEI turnover kinetic rate models.

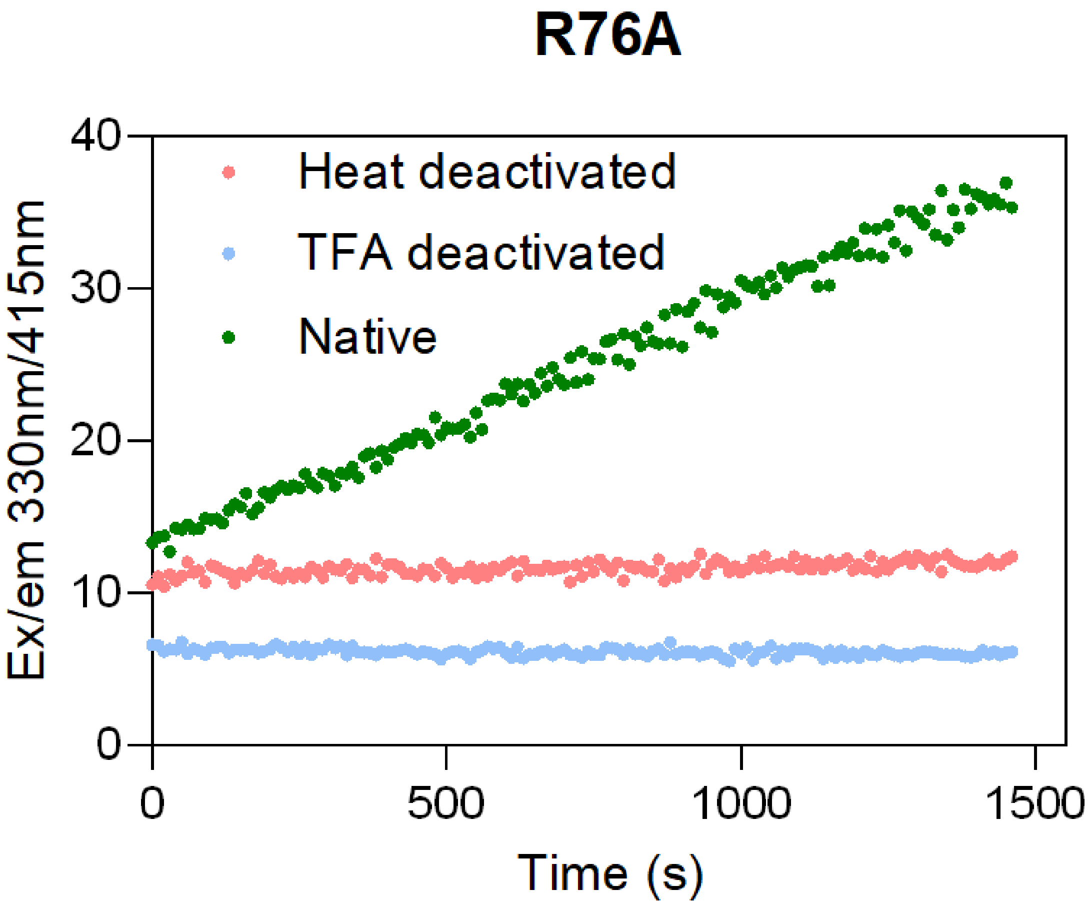


**Figure S4**. Validation of the Dark Mutant Activity. R76A was deactivated both through acid quenching and heat to corroborate inactivity of the enzyme, while stable R76A showed production of CEI over time under the signature CEI fluorescence excitation and emission wavelengths.

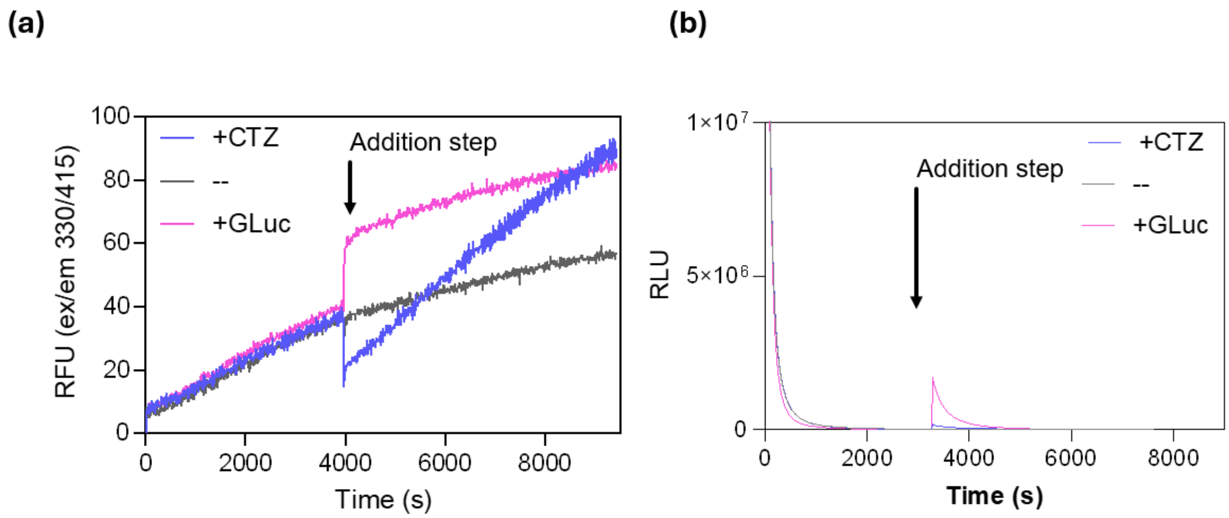


**Figure S5**. We assessed whether fresh substrate or fresh luciferase would replenish an ongoing reaction between GLuc and CTZ. CEI fluorescent monitoring in (a) reveals that the addition of either enzyme or substrate continued the production of CEI over time. However, luminescence monitoring in panel (b) shows that fresh enzyme replenishes luminescent signal much more effectively than fresh substrate.

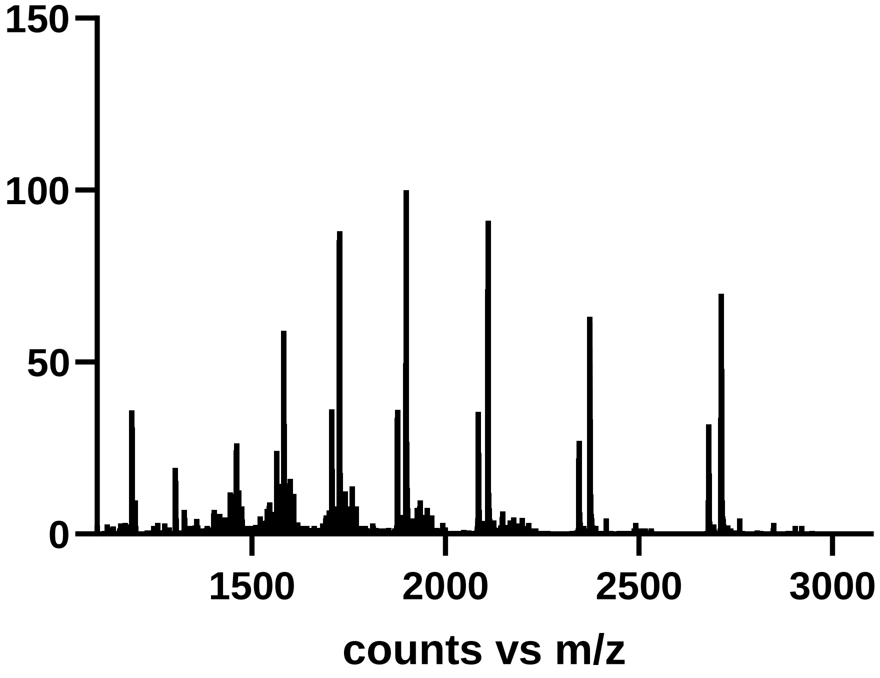


**Figure S6**. Raw Source and Deconvoluted ESI-Mass Spectrometry data of raw Gluc from the data presented in Figure 3 of the main manuscript. Raw spectra show signal was not compromised by salt or interfering detergents.

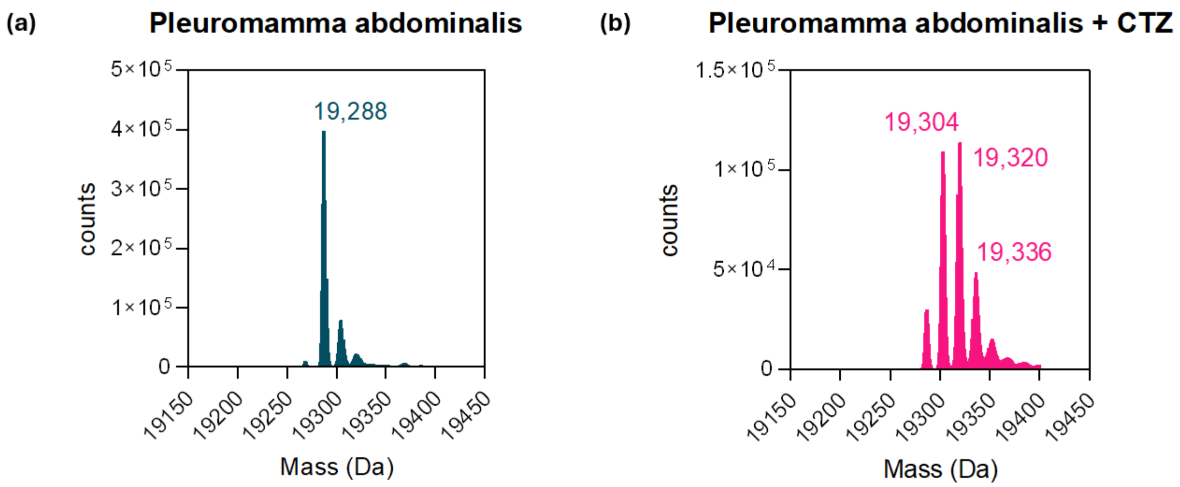


**Figure S7**. Deconvoluted ESI-Mass Spectrometry data of PaLuc2 and PaLuc2 reacted with CTZ shown in (a) and (b) respectively, reveal that an additional oxidation event occurs in PaLuc2, presumably by the additional methionine in the sequence. We believe that the subtle +16 and +32 peaks from the raw PaLuc2 sample are from nonspecific oxidation upon the protein during storage.

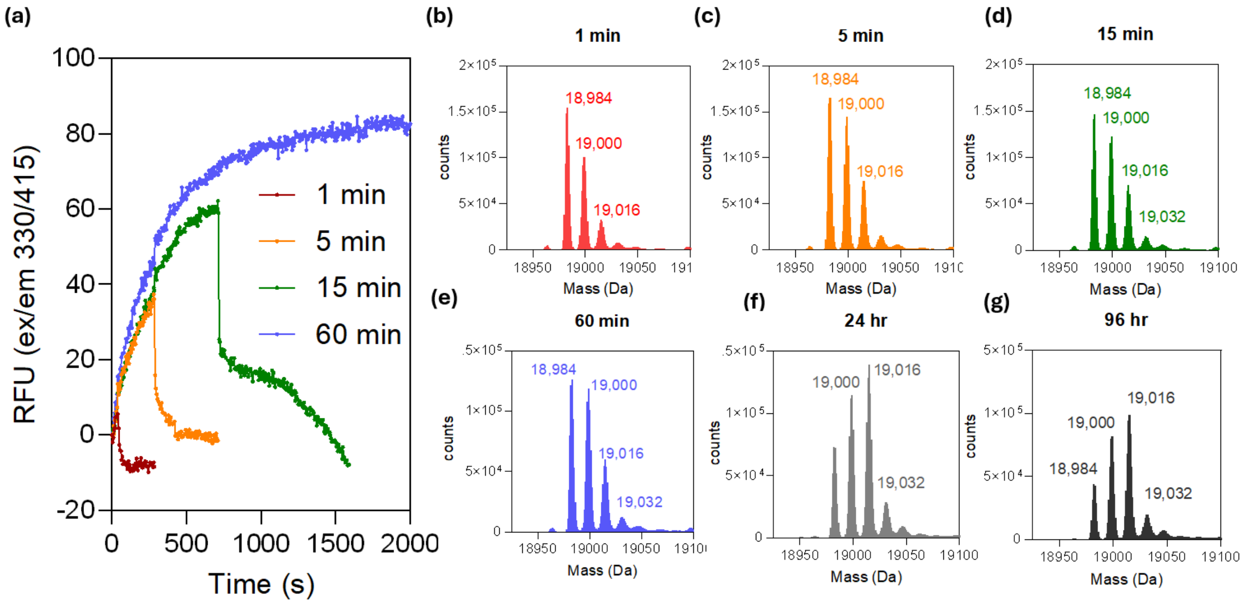


**Figure S8** Kinetic-based time point ESI-Mass spectrometry measurements of GLuc oxidized species after reaction with CTZ. The cessation of growth in fluorescent signal from the quenched reactions (1, 5, 15 min) can be seen and used as verification.

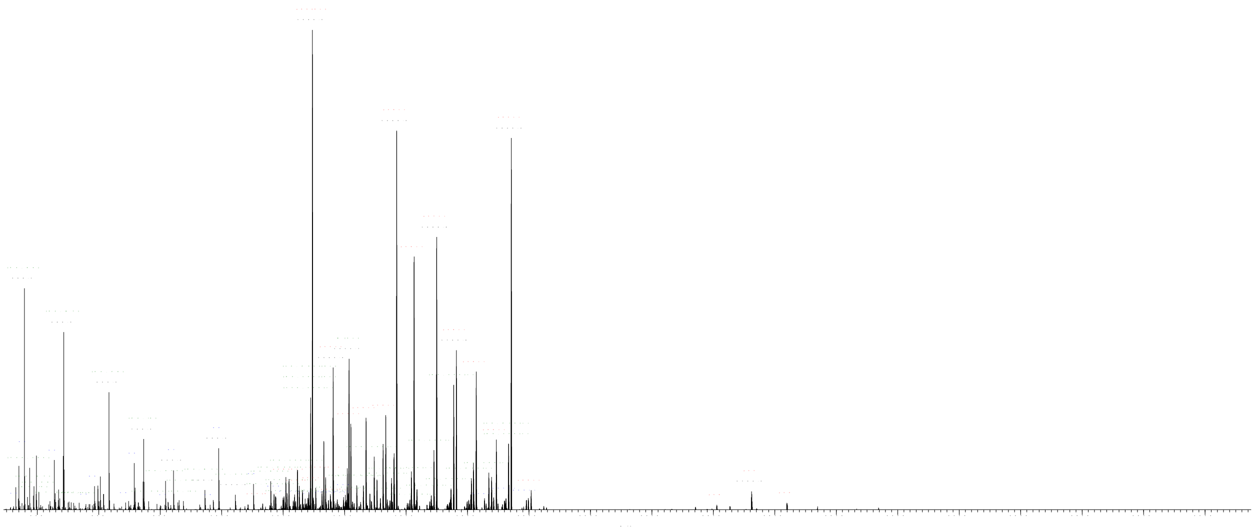

**Figure S9** – Annotated HCD MS2 spectrum of triply disulfide-bound L7-L13 peptides. Identified fragment ions corresponding to backbone cleavage of L13 including internal fragments are labelled in the spectrum. Generally, fragmentation is suppressed between and C120 and C127, consistent with the presence of disulfide bonds. Complete suppression of y-type fragmentation after C120 may suggest the presence of a C120-C127 disulfide bond, however the identity of the disulfide bond isoform cannot be identified uniquely.

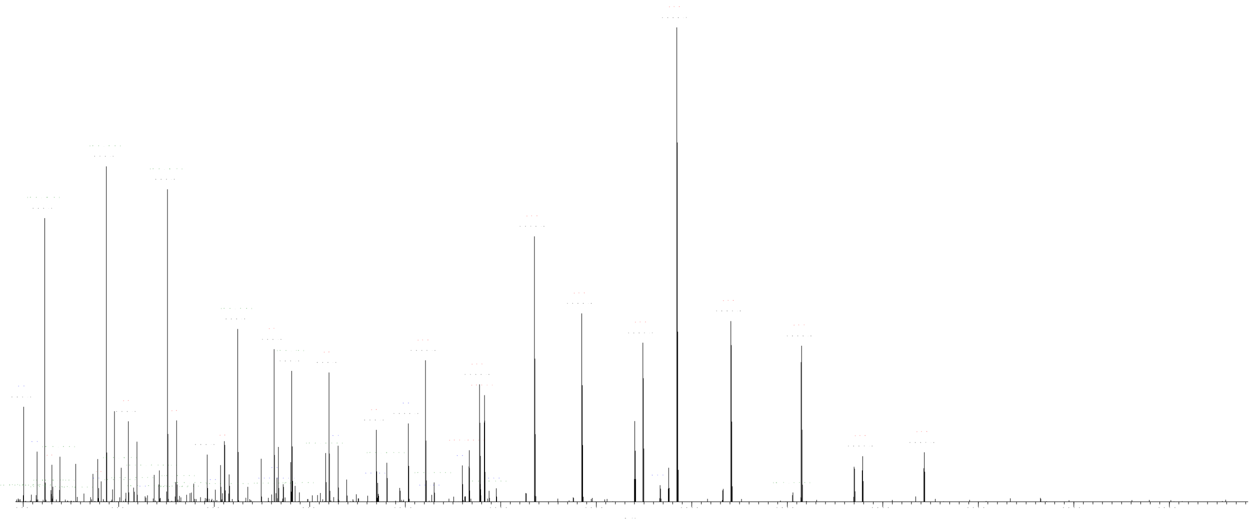

**Figure S10** – Annotated HCD MS2 spectrum of reduced peptide L13. Identified fragment ions are labelled in the spectrum. Near-complete y-type fragmentation is observed throughout the peptide backbone, alongside "-64" Dalton fragments diagnostic of methionine oxidation

Internal Use Only Ethics and Compliance

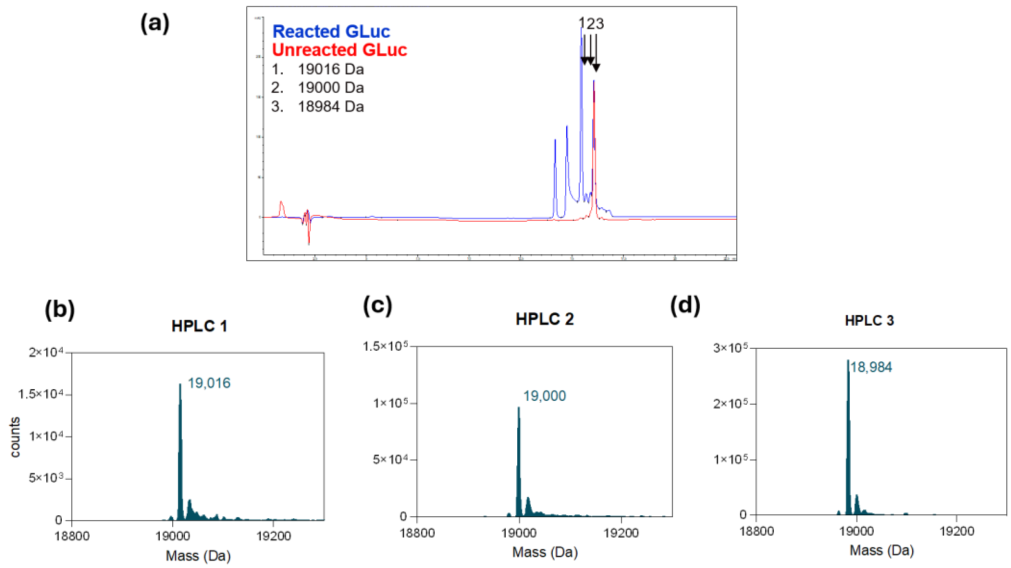


**Figure S11**. (a) show that both unreacted and reacted luciferase share a common peak "peak 3" of unreacted species, while additional peaks 1 and 2 arise from the reacted luciferase sample that are not present in the CTZ or CEI chromatograms. Peaks 1 and 2 correspond to the oxidized species (+32 and +16 respectively) from the reaction. Corresponding ESI Mass-Spectrometry data for the collected eluted fractions 3,2, and 1 for (a) and for GLuc demonstrate that the oxidized species can be separated from RP-HPLC

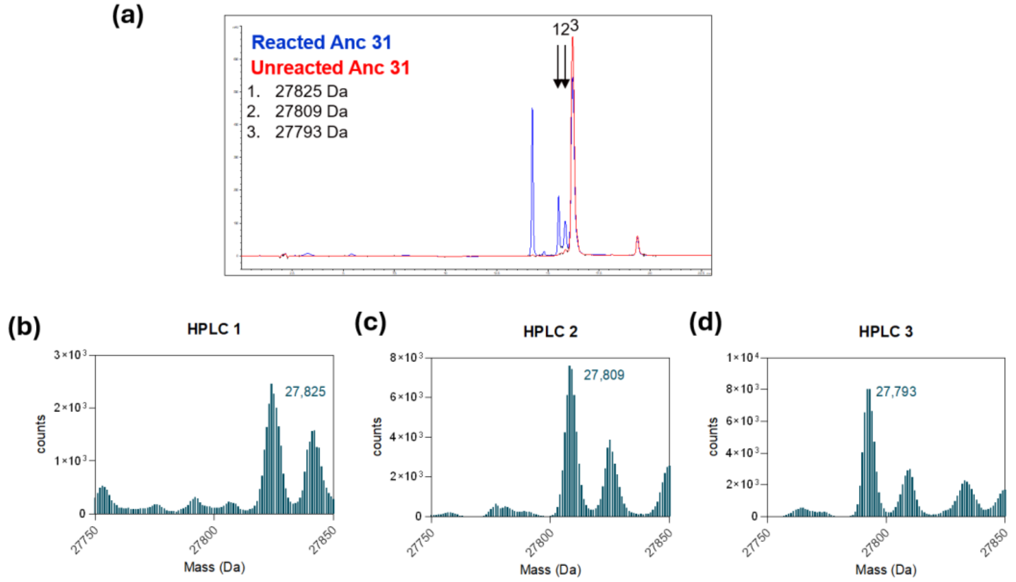


**Figure S12** (a) show that both unreacted and reacted luciferase share a common peak "peak 3" of unreacted species, while additional peaks 1 and 2 arise from the reacted luciferase sample that are not present in the CTZ or CEI chromatograms. Peaks 1 and 2 correspond to the oxidized species (+32 and +16 respectively) from the reaction. Corresponding ESI Mass-Spectrometry data for the collected eluted fractions 3,2, and 1 for (a) and for ANC 31 demonstrate that the oxidized species can be separated from RP-HPLC

GLuc WT

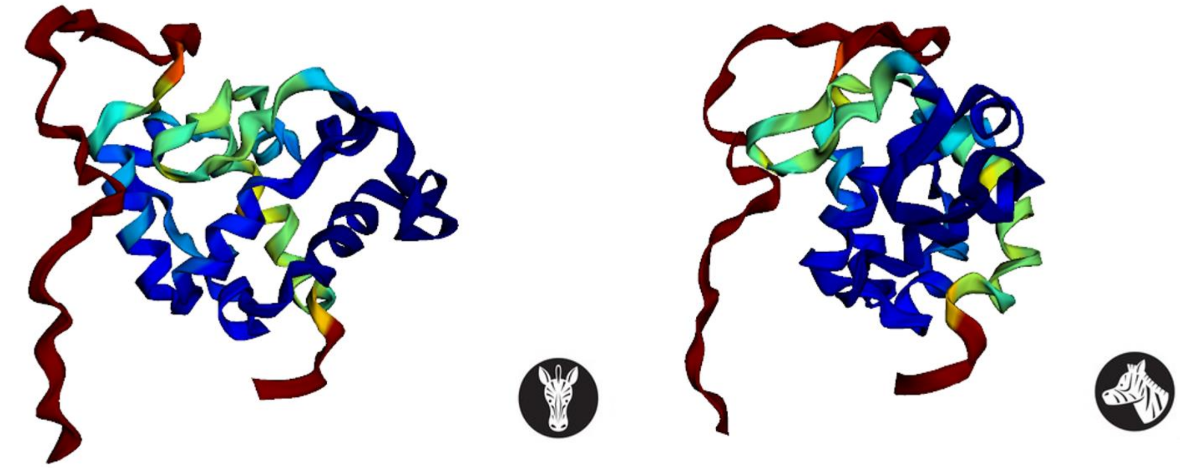

**Figure S13** representative AlphaFold2 predicted structure of Wildtype Gluc with no signal peptide

ANC 26

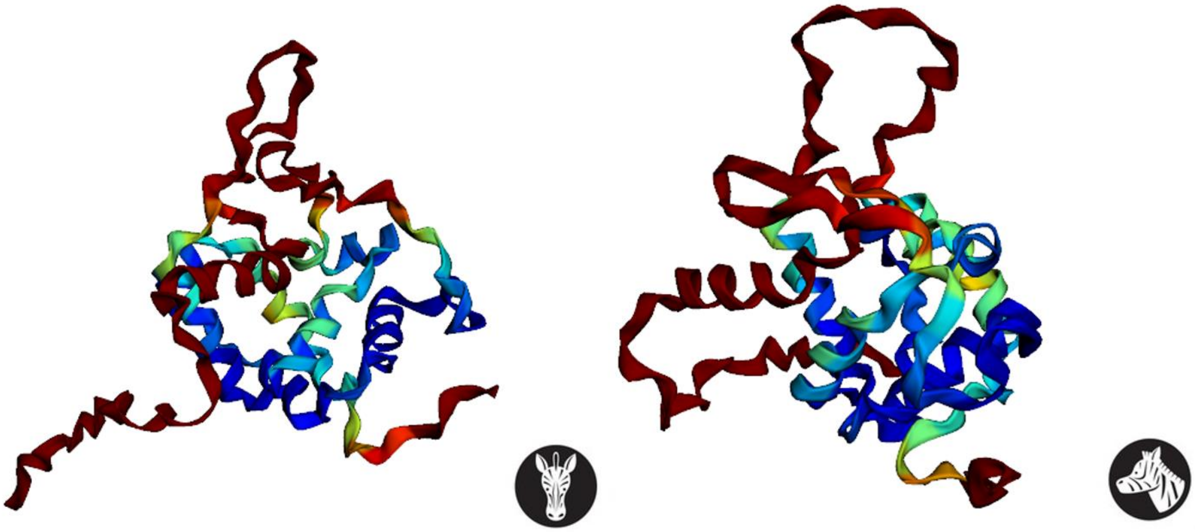

**Figure S14** representative AlphaFold2 predicted structure of ANC 26 with no signal peptide

Internal Use Only Ethics and Compliance

ANC 31

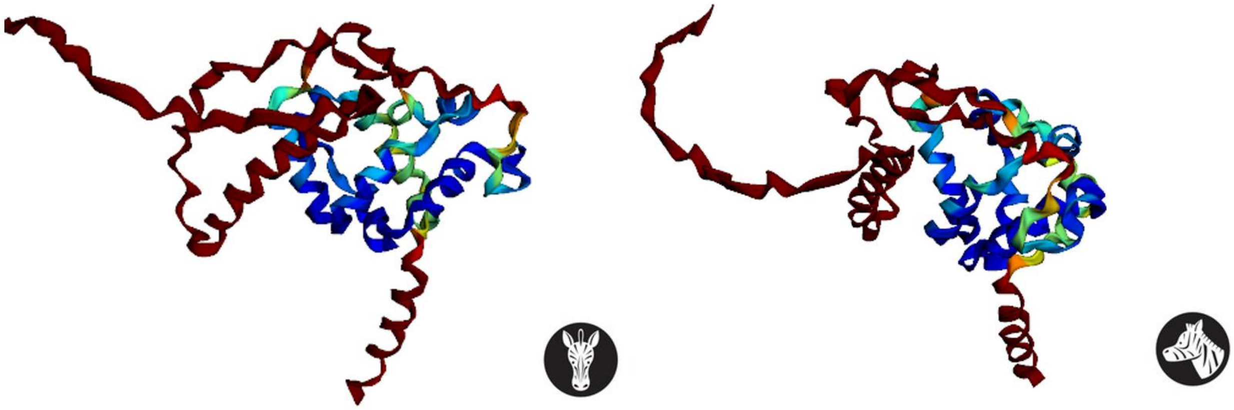

**Figure S15** representative AlphaFold2 predicted structure of ANC 31 with no signal peptide

ANC 35

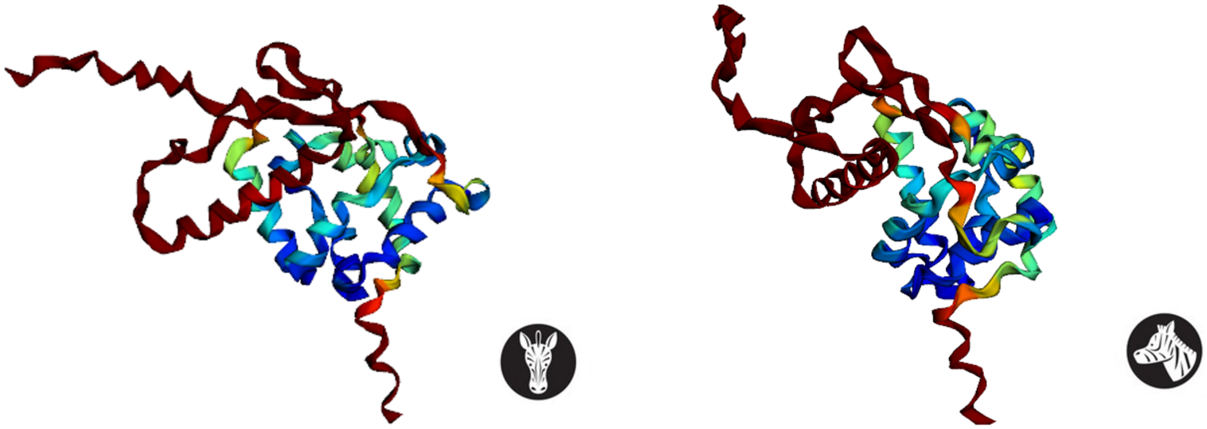

**Figure S16** representative AlphaFold2 predicted structure of ANC 35 with no signal peptide

Internal Use Only Ethics and Compliance

ANC 36

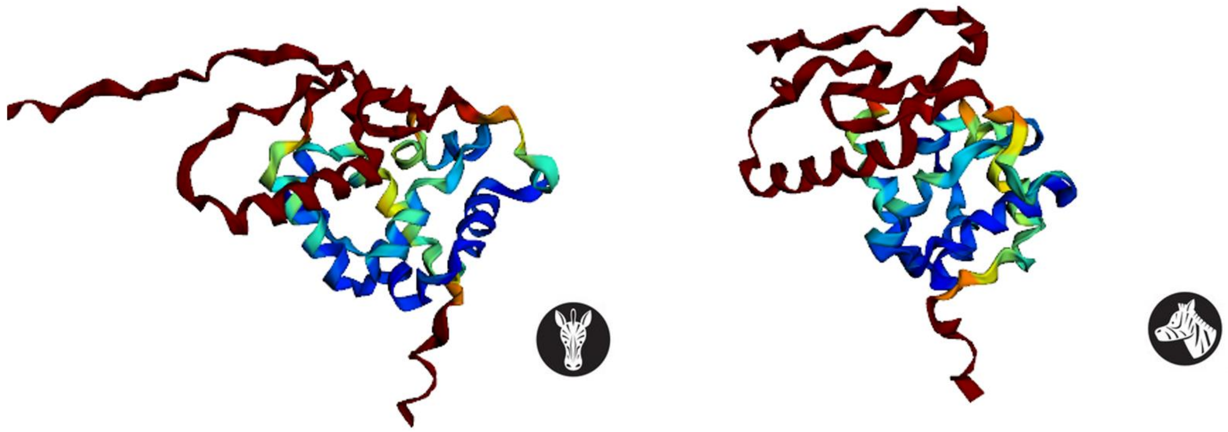

**Figure S17** representative AlphaFold2 predicted structure of ANC 36 with no signal peptide

ANC 37

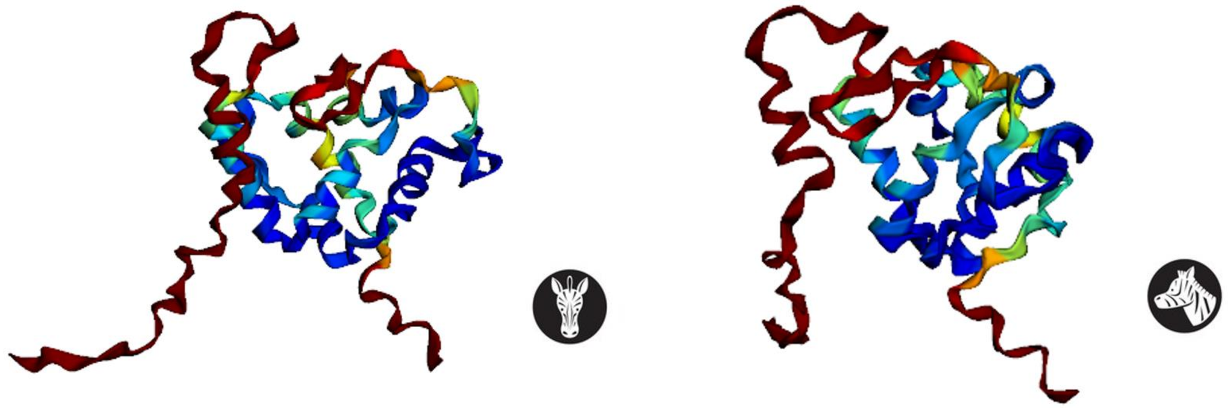

**Figure S18** representative AlphaFold2 predicted structure of ANC 37 with no signal peptide
